# HERC5-catalyzed ISGylation potentiates cGAS-mediated innate immunity

**DOI:** 10.1101/2023.01.03.522548

**Authors:** Lei Chu, Yu Chen, Li Qian, Wei Meng, Juanjuan Zhu, Quanyi Wang, Chen Wang, Shufang Cui

## Abstract

The key cytosolic DNA sensor cyclic GMP-AMP (cGAMP) synthetase (cGAS) is essential to elicit anti-infection innate immunity whereas aberrant activation of cGAS by endogenous DNA promotes severe autoimmune diseases, thus the activity of cGAS must be strictly regulated to maintain immune homeostasis. Here we report that the E3 ISG15-protein ligase HERC5 interacts with and catalyzes the cGAS ISGylation robustly, whereas the deconjugating enzyme USP18 removes the ISG15 conjugation of cGAS. The interaction of cGAS and HERC5 depends on the cGAS N-terminal domain and the RRC1-4 and HECT domain of HERC5. Mass spectrometry reveals that the Lys21,187,219,458 are the potential ISGylation sites in human cGAS. Deficiency of ISG15 or the E1 ubiquitin-activating enzyme UBA7 greatly attenuate the downstream gene expression induced by cGAS-STING axis and the antiviral ability in mouse and human cells. Collectively, our study uncovered an important role of the HERC5-facilitated cGAS ISGylaion in promoting antiviral defense.

## Introduction

The innate immune system employs germline-encoded pattern-recognition receptors to detect conserved pathogen associated molecular patterns (PAMPs) from pathogens, providing a first line of defense against pathogen invasion. Cytosolic aberrant DNA is generally sensed as a danger signal and evokes the innate immune DNA sensing pathway to control pathogen evasion. Cyclic GMP-AMP (cGAMP) synthase (cGAS), a recently defined key sensor of cytosolic double-strand DNA (dsDNA), triggers the type I interferon (IFN)-mediated innate immune signaling in a DNA-sequence-independent manner(1). Upon interacting with dsDNA, cGAS is activated and generates the second messenger 2’3’-cGAMP(2), which then binds to the endoplasmic reticulum protein stimulator of interferon genes (STING), ultimately inducing the expression of type I IFNs and interferon-stimulated genes (ISGs) in a TBK1-IRF3 axis-dependent manner(3, 4). Although indispensable for pathogen defense, inappropriate activation of cGAS by aberrant accumulated endogenous DNA is a key driver of autoimmune diseases such as Aicardi-Goutières syndrome (AGS) and familial chilblain lupus in human patients(5–7). Thus, the activity of cGAS must be tightly regulated spatially and temporally to maintain immune homeostasis.

Protein post-translational modifications (PTM) can shape the duration and strength of signal transduction effectively and quickly. Ubiquitylation and ubiquitylation-like modification such as sumoylation and neddylation, have been reported to regulate the DNA binding ability and nucleotidyl-transferase activities of cGAS(8, 9). Interferon-stimulated genes 15 (ISG15) is a ubiquitin-like protein which can be covalently conjugated onto target proteins by a three-step enzymatic cascade known as ISGylation. ISG15 and the members that mediate ISG15 conjugation can be induced by type I IFN rapidly and robustly(10). Briefly, the activating E1 enzyme UBE1L forms a thioester bond with the C-terminal glycine residue of ISG15(11). After activated, ISG15 is transferred to the active-site cysteine of the E2 ubiquitin/ISG15-conjugating enzyme UBE2L6(12). Then ISG15 is transferred to an E3 ligase which mediates the conjugation of ISG15 to the target proteins. UBE1L is the only identified E1 enzyme and functions exclusively in the ISG15 pathway. Whereas several E3 ligase have been identified in ISG15 conjugation, human HERC5 (mouse HERC6) is the dominant E3 ligase in the majority ISGylation(13, 14). Besides conjugated to target protein, ISG15 also exists non-covalently and functions as a soluble molecule with several immunomodulatory activities(15–17).

Recent studies demonstrate that ISG15 can limit viral infection by conjugated to both host and viral proteins. For example, HERC5 mediated IRF3 ISGylation resulted in sustained IRF3 activation, boosting the host antiviral immunity(18). The ISGylation of non-structural protein 1 of IAV (NS1/A) inhibited its nuclear translocation, resulting the virus susceptible to interferon(19). In contrast, viruses evolve different tactics to antagonize the ISGylation pathway, emphasizing the importance of this pathway in the host antiviral response. Recent study showed that the ISG15 conjugation of MDA5 was antagonized through direct de-ISGylation mediated by the papain-like protease of SARS-CoV-2, facilitating the immune-evasion of SARS-CoV-2(20). Although efforts have been done to clarify how ISG15 protects the host during infection, how ISG15 modulates the activity of cGAS is largely unknown. In this report, we found that cGAS interact specifically with the E3 ligase HERC5. Upon stimulation with IFN-β, cGAS co-localized with HERC5 and ISG15 robustly in the cytoplasm. Mechanically, HERC5 catalyzed the multi-ISGylation of cGAS, enhancing its antiviral ability. Knockdown of the E1 enzyme UBA7 or ISG15 crippled cGAS-mediated type I IFN expression and facilitated HSV-1 infection.

## Materials and methods

### Cell Culture and Transfection

HFF, HEK293T, HAEC, and Vero cells were cultured in DMEM medium (Invitrogen), supplemented with 10% FBS (Gibco) and 1% penicillin–streptomycin (Invitrogen). L929 cells were cultured in RMPI-1640 (Gibco) plus 10% FBS and 1% penicillin–streptomycin. All cells were cultured at 37°C and 5% CO2. Lipofectamine 3000 (Invitrogen) was used for transfection according to the manufacture’s procedure.

### Virus

HSV-1, GFP-HSV-1 were propagated and titrated by standard plaque assay on Vero cells.

### Reagents and Antibodies

HT-DNA were purchased from Sigma-Aldrich. Antibodies: anti-cGAS (CST, #31659), anti-GAPDH (Santa Cruz, sc-32233), anti-ISG15 (Santa Cruz, sc-166755), anti-UBA7(ABclonal, A9142), anti-UBE2L6(ABclonal, A13670), anti-HERC5(ABclonal, A14889), anti-Flag (Sigma, F3165), anti-Myc (ABclonal, AE010), anti-Rabbit IgG (H+L) (Jackson, 111-035-003), anti-MouseIgG (H+L) (Jackson, 115-035-003).

### Western Blot Analysis

Cell pellets were collected and lysed with lysis buffer (50 mM Tris-HCl pH 7.5, 150 mM NaCl, 1% Triton X-100) containing 1×EDTA-Free Protease inhibitor cocktail (Roche) and 1 mM PMSF (Sigma-Aldrich). The resuspended cell pellets were incubated on ice for 30 min, followed by centrifugation at 12,000 rpm for 15 min at 4°C. The supernatants were collected for SDS-PAGE. The bands were probed with indicated primary and secondary antibodies and visualized using a SuperSignal West Pico chemiluminescence ECL kit (Pierce).

For denaturing immunoprecipitation, cells were lysed in 1% SDS buffer (50 mM Tris-HCl pH 7.5, 150 mM NaCl, 1% SDS, 10 mM DTT) and denatured by heating for 30 minutes. The lysates were centrifuged and diluted with Lysis buffer (50mM Tris-HCl pH 7.5, 150 mM NaCl, 1 mM EDTA, 1% Triton X-100) until the concentration of SDS was decreased to 0.1%. The diluted lysates were immunoprecipitated with the indicated antibodies for 4 hours to overnight at 4°C before adding protein A/G agarose for 2 hours. After extensive wash, the immunoprecipitates were subjected to immunoblot analysis.

### Immunofluorescence and Confocal Microscopy

Cells were fixed for 15 min with 4% paraformaldehyde in PBS, washed with PBS, and permeabilized with 0.25% Triton X-100 in PBS for 20 min, then washed and blocked with 5% BSA for 1 h. After incubation with indicated primary antibodies overnight at 4°C, fluorescent-conjugated secondary antibodies were incubated for 2 h at room temperature. Slides were mounted with a fluorescent mounting medium (Dako) after DAPI (Sigma Aldrich) staining and captured using a confocal microscope (LSM700, Carl Zeiss) with a 63× oil objective.

### RNA Isolation and Quantitative PCR Analysis

Total RNA from cells was isolated with TRIzol reagent (Invitrogen) according to the manufacturer’s instruction. cDNA was synthesized with 0.5 ug RNA and HiScript III Q RT SuperMix for qPCR (Vazyme). Real-time PCR was performed using ChamQ SYBR qPCR Master Mix (Low ROX Premixed) (Vazyme). Data were normalized to the expression of mouse *Gapdh* or human *GAPDH*

through the comparative Ct method (2-ΔΔCt).

### Plaque Assay

The titers of the virus in the supernatant were determined by standard plaque assay with Vero cells.

After infection with virus contained supernatant for 2 h, cell monolayers were washed twice with pre-warmed DMEM medium, followed by incubation with a mixed media (1× MEM, 2% FBS, 1% Low Melting Point Agarose) until the formation of the plaques. Then the cells were fixed with 4% paraformaldehyde and stained with crystal violet.

### Statistical Analyses

Statistical analyses were performed with GraphPad Prism 8. A standard two-tailed unpaired Student’s t-test was performed for statistical analyses of two groups or as indicated in the figure legends. All analyzed data are expressed as mean ± standard deviation (SD). Differences with a P-value <0.05 were considered significant.

## Results

### cGAS interacts with the ISGylation E3 ligase

To explore whether ISGylation modulates the cGAS-STING pathway, we first examined the interaction of ISGylation E3 ligase with cGAS. The immunoprecipitation (IP) results showed that the dominate E3 ligase HERC5 interacted robustly with cGAS, but not STING (Figure 1A). Consistent with the previous study, there was a strong interaction of HERC5 and IRF3 (Figure 1A). In addition, we observed a weakly interaction of cGAS with EFP (TRIM25) (Figure 1B). Then we checked which domains of each protein mediated their interaction by expressing cDNA fragments encoding the different truncations of cGAS and HERC5 in HEK293T cells. The IP results showed that the N-terminal domain of cGAS (aa 1-160) was required for its interaction with HERC5 (Figure 1C). We found that HERC5 interacted with cGAS dependent of two domains, the RRC1-4 (aa 262-312) and the HECT domain (aa 702-1024) (Figure 1D), which is responsible for interact with a ubiquitin-conjugating enzyme(13). Taken together, these data demonstrate that cGAS interacts with the ISGylation E3 ligase, and cGAS might be a potential candidate of ISGylation.

**Figure 1.**
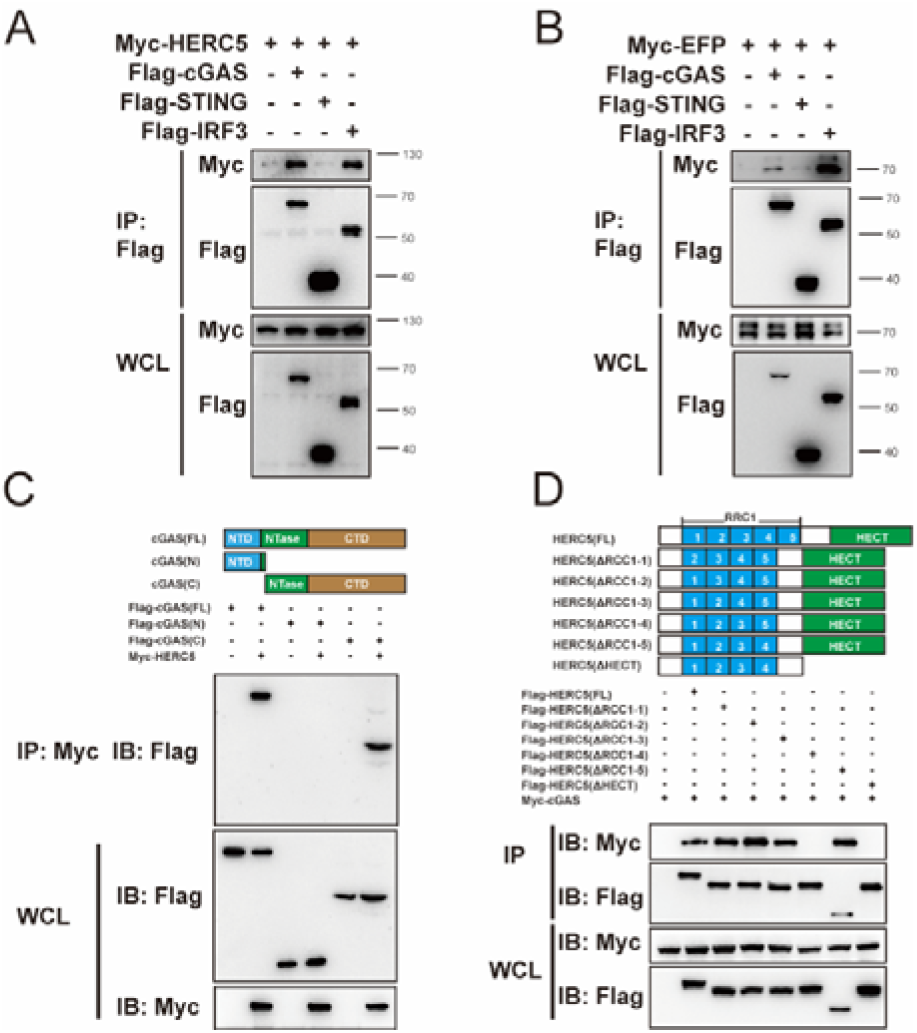
cGAS interacts with the ISGylation E3 ligase. A. Immunoprecipitation (with anti-Flag) and immunoblot analysis (with anti-Flag or anti-Myc) in HEK293T cells that were transfected with plasmids encoding Myc-HERC5 and Flag-tagged, cGAS, STING or IRF3. B. Immunoprecipitation (with anti-Flag) and immunoblot analysis (with anti-Flag or anti-Myc) in HEK293T cells that were transfected with plasmids encoding Myc-EFP and Flag-tagged, cGAS, STING or IRF3. C. Immunoprecipitation (with anti-Myc) and immunoblot analysis (with anti-Flag or anti-Myc) in HEK293T cells that were transfected with plasmids encoding Myc-HERC5 and Flag-tagged cGAS or truncates. NTD N-terminal domain, NTase nucleotidyl transferase, CTD C-terminal domain. D. Immunoprecipitation (with anti-Flag) and immunoblot analysis (with anti-Flag or anti-Myc) in HEK293T cells that were transfected with plasmids encoding Myc-cGAS and Flag-tagged HERC5 or truncates. RCC1 regulator of chromosome condensation 1, HECT homologous to the E6AP carboxyl terminus. All data shown are representative of three independent experiments.

### HERC5 catalyzes the multi-ISGylation of cGAS

As cGAS can interact with the E3 ligase, next we investigated whether cGAS could be ISGylated. Exogenous ISG15, cGAS and the ISGylation machinery (including UBA7, UBE2L6, HERC5 or EFP) were transfected into HEK293T cells followed by denatured-immunoprecipitation assay with Ni-NTA beads. The result showed that cGAS was multi-ISGylated by HERC5 but not EFP (Figure 2. A-B). In addition, we found that the ISGylation of cGAS could be reversed by the deconjugating enzyme Ubl carboxy-terminal hydrolase 18 (USP18) (Figure 2B), a specific ISG15 deconjugase (21). To further confirm the HERC5-catalyzed cGAS ISGylation, we repeated the denatured-immunoprecipitation experiment with Flag beads and obtained consistent result (Figure 2C). In line with the previous report and the results in Figure 1, we found that HERC5 could catalyze the ISGylation of cGAS and IRF3, rather than STING (Figure 2D). Since the conserved cysteine 994 residue residing in the HECT domain of HERC5 is crucially important for its ISG15 E3 ligase activity, next we went on to address whether this site was indispensable for mediating cGAS ISGylation(13). As expected, the enzymatic dead mutant HERC5 C994A lost its ability to catalyze ISGylation of cGAS (Figure 2E). As ISG15 is covalently conjugated to the target proteins through its carboxy-terminal LRLRGG motif, we constructed cDNA fragments encoding the mutant ISG15 (RGG to RAA) and expressed them in HEK293T cells together with cGAS and the ISGylation machinery. The result showed that cGAS could be ISGylated by wild-type ISG15 but not the ISG15 plasmid lost its covalent ability (Figure 2F). Taken together, these results demonstrate that cGAS can be ISGylated specifically catalyzed by HERC5.

**Figure 2.**
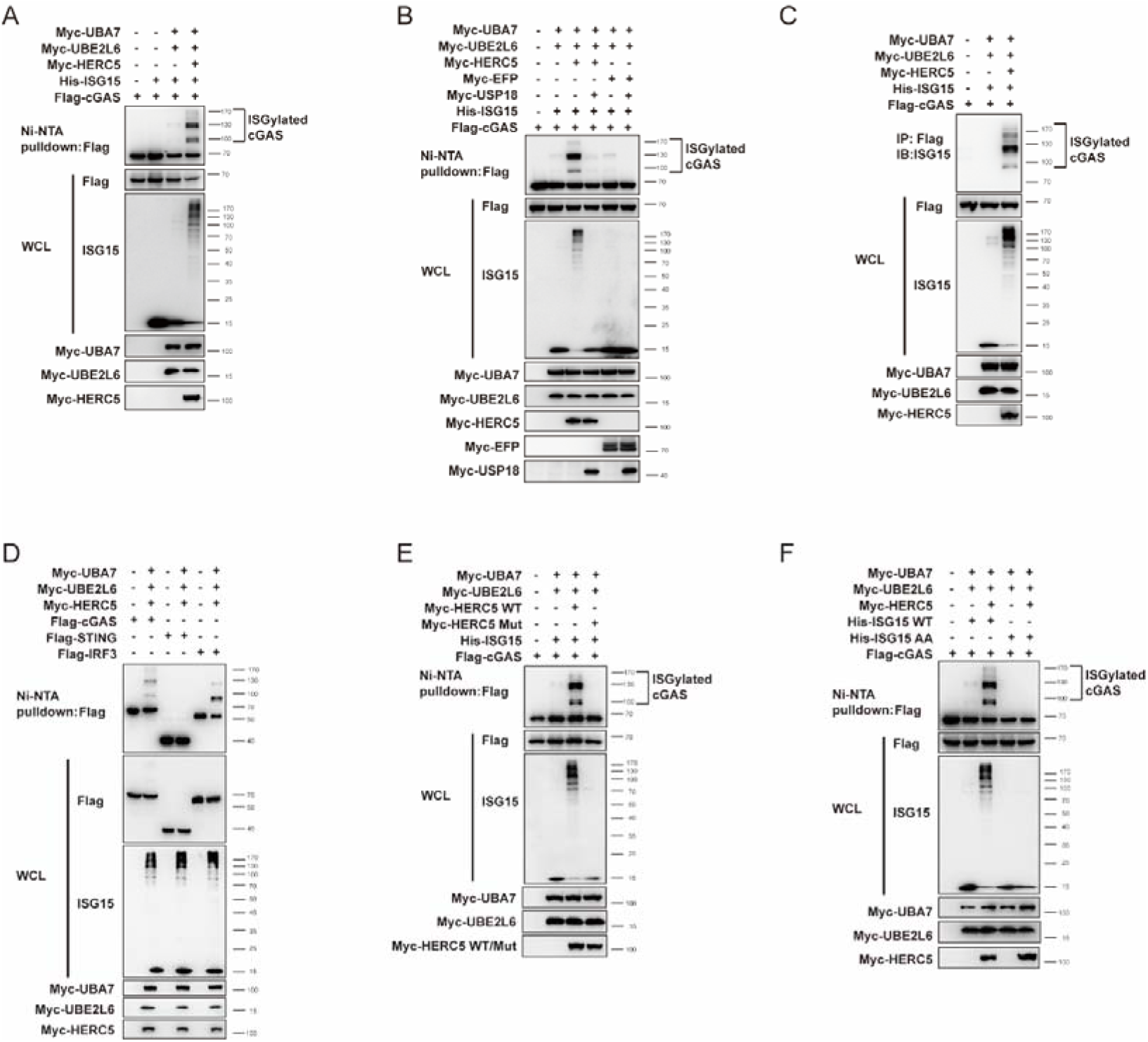
HERC5 catalyzes the multi-ISGylation of cGAS. A. Denature-IP (Ni-NTA pulldown) and immunoblot analysis (with anti-Flag, anti-Myc, or anti ISG15) of HEK293T cells transfected with plasmids encoding Flag-cGAS, His-ISG15, and Myc-tagged UBA7, UBE2L6 or HERC5. B. Denature-IP (Ni-NTA pulldown) and immunoblot analysis (with anti-Flag, anti-Myc, or anti ISG15) of HEK293T cells transfected with plasmids encoding Flag-cGAS, His-ISG15, and Myc-tagged UBA7, UBE2L6, HERC5, EFP or USP18. C. Denature-IP (with anti-Flag) and immunoblot analysis (with anti-Flag, anti-Myc, or anti ISG15) of HEK293T cells transfected with plasmids encoding Flag-cGAS, Flag-STING, Flag-IRF3, His-ISG15, and Myc-tagged UBA7, UBE2L6 or HERC5. D. Denature-IP (Ni-NTA pulldown) and immunoblot analysis (with anti-Flag, anti-Myc, or anti ISG15) of HEK293T cells transfected with plasmids encoding Flag-cGAS, His-ISG15, and Myc-tagged UBA7, UBE2L6 or HERC5. E. Denature-IP (Ni-NTA pulldown) and immunoblot analysis (with anti-Flag, anti-Myc, or anti ISG15) of HEK293T cells transfected with plasmids encoding Flag-cGAS, His-ISG15, and Myc-tagged UBA7, UBE2L6, HERC5 WT or HERC5 C994A. F. Denature-IP (Ni-NTA pulldown) and immunoblot analysis (with anti-Flag, anti-Myc, or anti ISG15) of HEK293T cells transfected with plasmids encoding Flag-cGAS, His-ISG15 WT, His-ISG15 AA and Myc-tagged UBA7, UBE2L6 or HERC5. All data shown are representative of three independent experiments.

### cGAS co-localize with HERC5 or ISG15 in the cytoplasm

In the cytosol, cGAS acts a canonical DNA sensor initiating downstream of immune signaling cascades. We next found that the predominant presence of ISGylated substrates in cytosol. Whereas cGAS overexpression showed its random distribution in cytoplasm or nucleus, only the cytosolic cGAS can be ISGylated and compose to cGAS-ISG15 puncta (Figure 3A). Notably, cGAS exhibited highly co-location with the two ISGs HERC5 and ISG15 after interferon ß stimulation (Figure 3C, E). Consistently, interferon-induced HERC5 endogenously associated with cGAS (Figure 3B). Moreover, we observed endogenous ISGylation of cGAS (Figure 3D). These data together suggest that cGAS interacts with HERC5 and can be ISGylated after interferon β stimulation.

**Figure 3.**
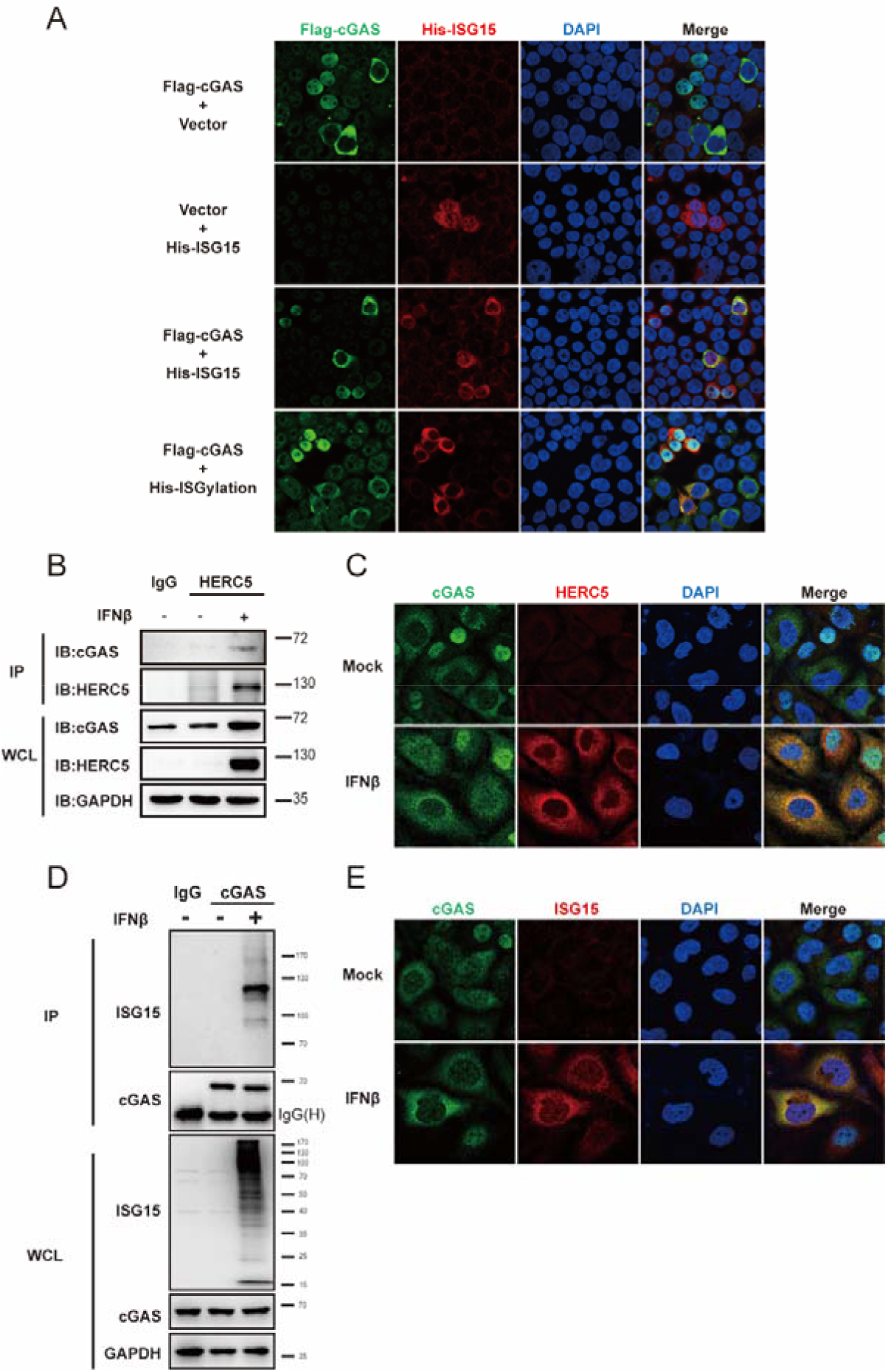
cGAS co-localize with HERC5 or ISG15 in the cytoplasm. A. Confocal microscopy of HEK293T cells transfected with plasmids encoding Flag-cGAS, His-ISG15, and Myc-tagged UBA7, UBE2L6 or HERC5. B. Immunoprecipitation (with control IgG or anti-HERC5) and immunoblot analysis (with anti-cGAS, anti-HERC5, or anti-GAPDH) in HAEC cell treated with or without interferon ß. C. Confocal microscopy of HAEC cell treated with or without interferon β, stained with indicated antibodies (anti-cGAS and anti-HERC5). D. Denature-IP (with control IgG or anti-cGAS) and immunoblot analysis (with anti-cGAS, anti-ISG15, or anti-GAPDH) in HAEC cell treated with or without interferon ß. E. Confocal microscopy of HAEC cell treated with or without interferon β, stained with indicated antibodies (anti-cGAS and anti-ISG15). All data shown are representative of three independent experiments.

### Crippled ISGylation decrease DNA-induced genes expression

It has been reported that UBA7/UBE1L is also an IFN-stimulated gene and function as only E1 for ISG15 conjugation(22). To explore whether ISGylation system regulated DNA-sensing signaling, we used two pairs of small interfering RNA (siRNA) to silence UBA7 and ISG15 in mouse L929 and human HFF cells. Both the siRNAs efficiently exhibited appropriate knockdown effects (Figure 4A-D). Quantitative analysis revealed that both silencing of Uba7 and Isg15 drastically reduced the expression of interferon and other cytokines (*Ifnb, Ifna4, Il6* and *Mcp-1*) in L929 upon HT-DNA stimulation (Figure 4A, B). Consistently, we also observed the suppression of ISGylation system leads to a significant decrease in the amounts of DNA-triggered antiviral genes expression in human HFF cells (Figure 4C, D). Together, these results demonstrate that ISGylation plays a positive role in regulating DNA-mediated innate immunity.

**Figure 4.**
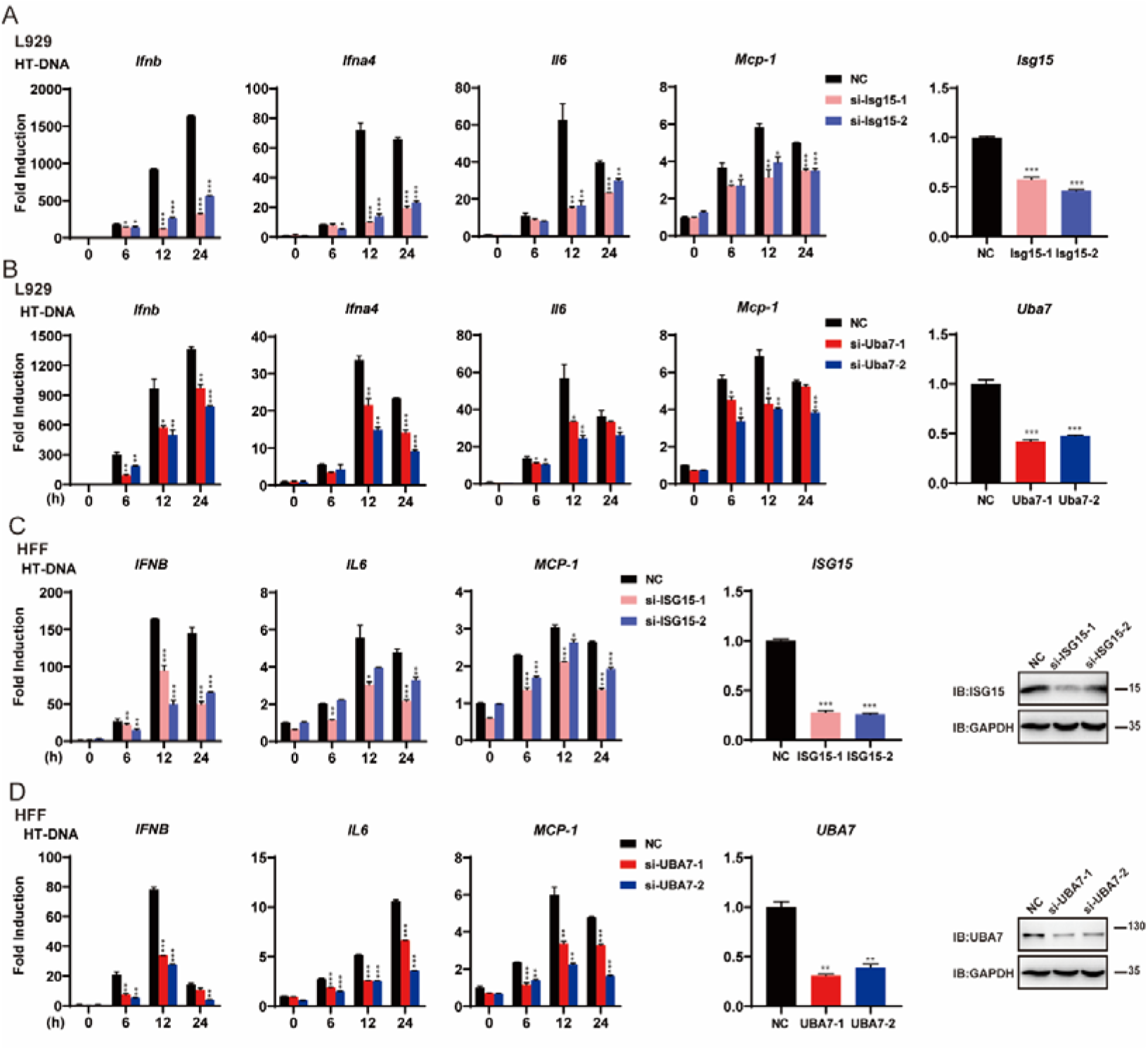
Crippled ISGylation decrease DNA-induced genes expression. A. L929 cells transfected with the nonspecific control (NC) or Isg15 siRNAs were stimulated with HT-DNA. Induction of *Ifnb, Ifna4, Il6, Mcp-1*and *Isg15* mRNAs were measured by quantitative PCR. B. L929 cells transfected with the nonspecific control (NC) or Uba7 siRNAs were stimulated with HT-DNA. Induction of *Ifnb, Ifna4, Il6, Mcp-1and Uba7* mRNAs were measured by quantitative PCR. C. HFF cells transfected with the nonspecific control (NC) or ISG15 siRNAs were stimulated with HT-DNA. Induction of *IFNB, IL6, MCP-1and ISG15* mRNAs were measured by quantitative PCR. The immunoblots (right panel) showed the knockdown of ISG15. D. HFF cells transfected with the nonspecific control (NC) or UBA7 siRNAs were stimulated with HT-DNA. Induction of *IFNB, IL6, MCP-1* and *UBA7* mRNAs were measured by quantitative PCR. The immunoblots (right panel) showed the knockdown of UBA7. Graphs show the mean ± s.d. and data shown are representative of three independent experiments. *P <0.05; **P <0.01; ***P <0.001 (two-tailed t-test).

### Inhibition of ISGylation promotes HSV-1 infection *in vitro*

Since cGAS-mediated immune responses protect host cells against viruses. To investigate the *in vitro* antiviral activity of ISGylation, we employed ISGylation-knockdown L929s before challenge with DNA viruses Herpes simplex virus 1 (HSV-1). Plaque assays indicated that both Isg15- and Uba7-kockdown increased the virus titers of the cell supernatants (Figure 5A). We also detected virus proliferation post-infection with GFP-expressing HSV-1. Similarly, knockdown of ISGylation increased the number of GFP-HSV-1-positive cells (Figure 5B). Collectively, these results confirm the positive role of ISGylation in regulating the innate immune response to DNA viruses *in vitro*.

**Figure 5.**
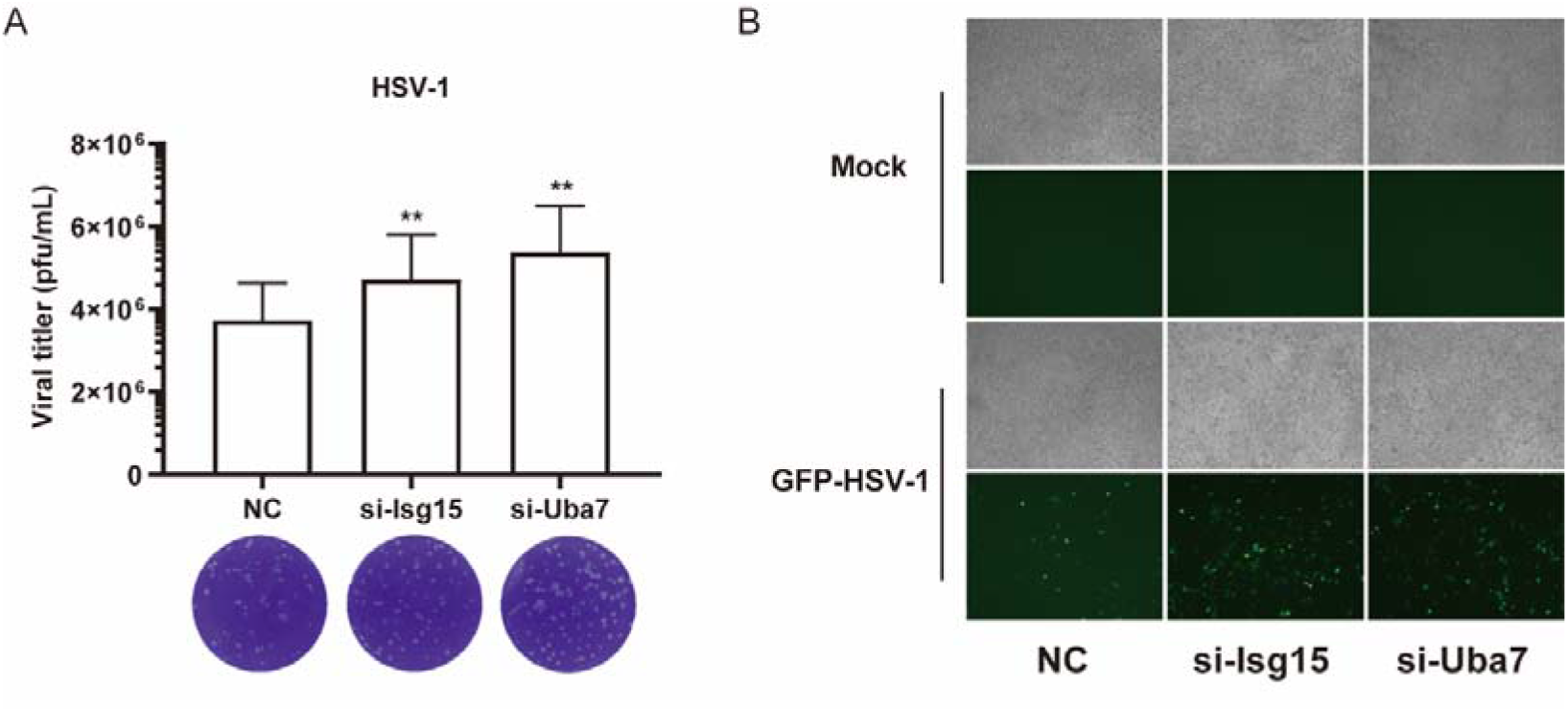
Inhibition of ISGylation promotes HSV-1 infection *in vitro*. A. L929 cells transfected with the indicated siRNAs were infected with HSV-1. The titers of HSV-1 were determined by standard plaque assay. B. GFP-HSV-1 replication in L929 cells transfected with the indicated siRNAs were visualized by fluorescence microscopy. Graphs show the mean ± s.d. and data shown are representative of three independent experiments. *P <0.05; **P <0.01; ***P <0.001 (two-tailed t-test).

### cGAS is multi-ISGylated at Lys21,187,219 and 458

Since Ub- or Ubl- conjugation to substrates depends on their C-terminal motif, conventional MS approach to identify ubiquitinated sites cannot discriminate the K-εGG peptides from the different modifications(23). To further identify which lysine residues on cGAS are modified by ISGylation, we performed a modified liquid chromatography-mass spectrometry analysis as shown in Fig 6A. We employed a “KGG” ISG15 mutant in overexpression ISGylation system, the products were digested only by Lys-C protease after the immunoaffinity purification. Thus, K-εGG peptides were generated specifically from ISG15 KGG-modified substrates. Next, we identified K21,187,219 and 458 as cGAS ISGylation sites with high confidence. Compared to cGAS WT, several K-to-R mutants had drastically reduced ISGylation signals in HEK293T cells (Figure 6B). These data demonstrate that cGAS can be multi-ISGylated on Lys21,187,219, and 458, respectively.

**Figure 6.**
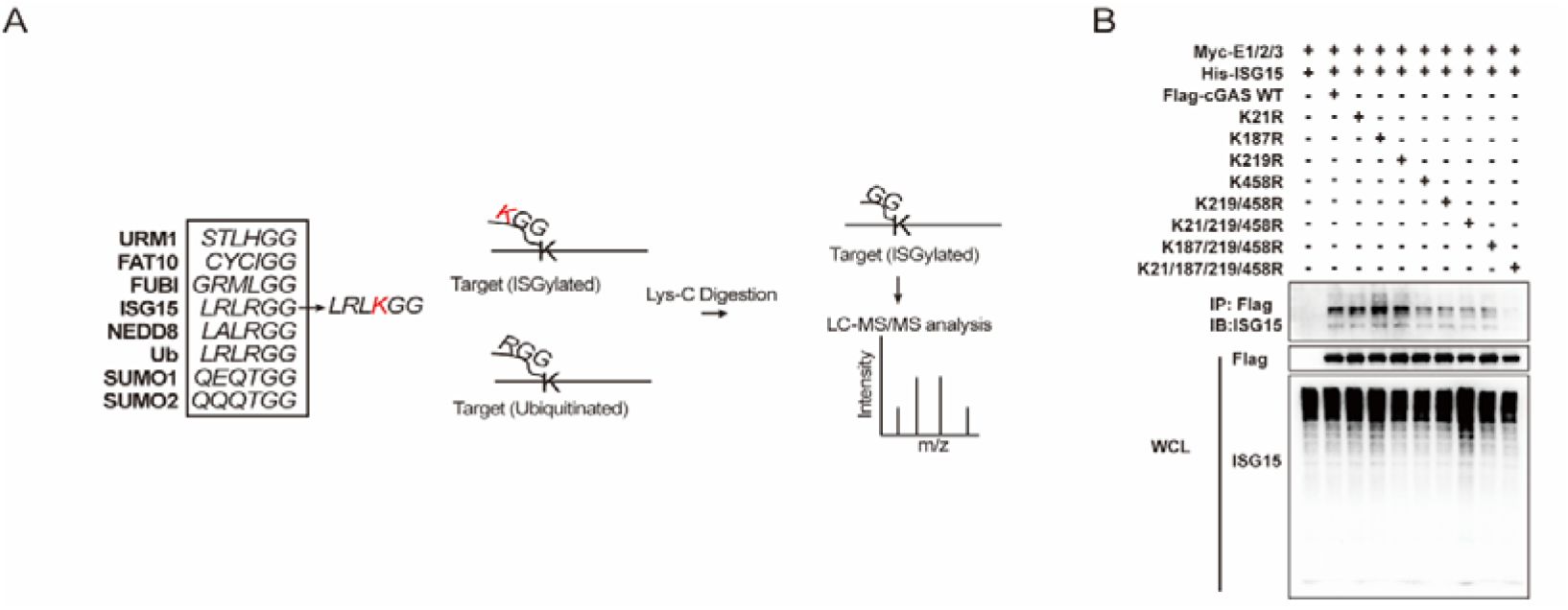
cGAS is multi-ISGylated at Lys21,187,219 and 458. A. A scheme showing modified liquid chromatography-mass spectrometry analysis for identify ISGylation sites. B. Denature-IP (with anti-Flag) and immunoblot analysis (with anti-Flag or anti ISG15) of HEK293T cells transfected with plasmids encoding Flag-cGAS WT or Mutants together with Myc-tagged UBA7, UBE2L6 andHERC5. All data shown are representative of three independent experiments.

## Discussion

ISG15 exists as ubiquitin-like protein that is rapidly induced to confer antiviral activity. Notably, the inductions of ISG15 conjugation cascades (UBA7, UBE2L6, HERC5, ISG15 and USP18) specially responses to ISRE elements through transcription factors IRFs(24). Thus, it’s not surprising that the unique PTM regulate the outcome of infection and pathogenesis. Aberrant-exposed DNA robustly initiates the rapid propagation of cGAS-mediated immune responses, which render the increase of overall ISGylation level. In turn, whether ISGylation may involves in the upstream of signaling sensing is not fully understood.

In this study, we identified that HERC5 interacts with cGAS robustly. As a canonical E3 ISG15–protein ligase, HERC5 specifically catalyze the multiple-sites ISGylation of cGAS via its enzymatic activity, but not STING. Notably, another substrate-specific E3 EFP cannot ISGylate cGAS, as well as USP18 removes the conjugated ISG15 from cGAS. Furthermore, cytosol cGAS but not nuclear cGAS undergoes ISGylation after interferon stimulation. Meanwhile, ISGylation is indispensable in DNA-induced antiviral responses. Inhibition of ISGylation may facilitate virus proliferation through dampen innate immune responses. Further modified proteomic studies reveled that cGAS could be ISGylated at 4 lysine residues.

Recent studies have suggested that ISGylation involve the nucleic acid-sensing pathways(18, 20, 25–27). As the major RNA-sensing RLRs members, both RIG-I and MDA5 are ISGylated upon virus infection. Moreover, the noncovalent form of ISG15 negatively regulating RIG-I-mediated signaling transduction through delivery RIG-I to autophagosome degradation. ISG15 and ISGylation function as potent antiviral elements in context of infections(28–31). Previous studies have shown that both host and viral proteins interacts with ISG15 which can disrupt viral replication(19, 32–34). Conversely, virus have adopted diverse strategies to counteract for ISGylation(20, 22, 35). SARS-CoV-2 encodes for a papain-like protease PLpro, which is a putative deISGylase facilitating immune evasion(35). The complexity of the regulatory networks related to ISG15 or ISGylation seems to far exceed the complexity of the immune system– activation mechanism itself, studies of these mechanisms in detail may provide a spectrum of beneficial consequences for therapeutic application.

In summary, our results uncover the function of ISGylation in the regulation of cGAS mediated type I IFN signaling. More importantly, we provide new insight that DNA-induced ISGylation act as a new PTM of cGAS and a new positive feedback mechanism of the innate immune response. Furthermore, since cGAS involves in many diseases, our findings have identified a novel therapeutic target for developing infection, inflammatory, and cancer therapies.

